## Supplementary material for "Species and condition shape the mutational spectrum in experimentally evolved biofilms": Table S1 to S3 and Figure S

### Table S1 Laboratory evolution setups of four experiments

| **Group** | **Bth_bead** | **Bth_root** | **Bs_pellicle** | **Bs_root** |
|---|---|---|---|---|
| Species | *B. thuringiensis* | *B. thuringiensis* | *B. subtilis* | *B. subtilis* |
| Adaptation condition | Nylon beads floating in the medium | *A. thaliana* root | Pellicle biofilm at the air-medium interface | *A. thaliana* root |
| Medium | EPS medium | MSNg medium | MSgg medium | MSNg medium |
| Medium volume | 1000 µL | 300 µL | 2000 µL | 300 µL |
| Temperature | 30C | 16 h light at 24 °C/8 h dark at 20 °C | 30C | 16 h light at 24 °C/8 h dark at 20 °C |
| Transfer interval time | 24h | 48h | 48h | 48h |
| Number of transfers | 40 | 38 | 35 | 32 |
| Number of parallel linages | 5 | 6 | 5 | 7 |
| Timepoints | 7 | 6 | 7 | 5 |
| Total population samples sequenced | 34 | 35 | 35 | 34 |
| Adaptation model | one colonized bead to **two** new beads | one colonized root to one new root | static floating biofilm disrupted by glass beads and vortexing for 1:100 reinoculation | one colonized root to one new root |
| Shaking condition | 90 rpm | 90 rpm | static | 90 rpm |

### Table S2 *dN/dS* ratio

| **Replicate** | **Bth_bead** | **Bth_root** | **Bs_pellicle** | **Bs_root** |
|---|---|---|---|---|
| R1 | 2.98 | 2.85 | 0.50 | 0.79 |
| R2 | * | 1.14 | 1.57 | 0.91 |
| R3 | 0.68 | 0.88 | 2.85 | 0.57 |
| R4 | 0.45 | 0.75 | 0.47 | 0.82 |
| R5 | 2.44 | 2.37 | 1.14 | 1.06 |
| R6 | 1.08 | | 0.78 | 1.99 |
| R7 | | | | 0.93 |
| Total ratio | 1.36 | 1.25 | 0.95 | 0.91 |
| Average | 1.53 | 1.60 | 1.22 | 1.01 |
| SD | 1.00 | 0.85 | 0.82 | 0.43 |

* no synonymous mutation detected in this lineage

**Table S3**

**A/ Transposase gene in the *B. thuringiensis* 407 genome**

| **Transposase gene** | **Copies** |
|---|---|
| IS*110* family transposase | 12 |
| IS*110*-like element ISBth13 family transposase | 5 |
| IS*21*-like element IS*232* family transposase | 5 |
| IS*3* family transposase | 5 |
| transposase | 5 |
| IS*4* family transposase | 2 |
| IS*4*-like element IS*231*A family transposase | 2 |
| IS*4*-like element IS*231*C family transposase | 1 |
| IS*607* family transposase | 1 |
| IS*66* family transposase | 1 |

**B/ Insertion sequence definition in the second-round analysis using *breseq***

| IS no. | Position | Length | Tag | Gene name |
|---|---|---|---|---|
| IS1 | 1903509~1905263 | 1755 | BTB_RS09780 | IS*4* like element IS*231*A family transposase |
| IS2 | 2497927~2499656 | 1730 | BTB_RS12605 | IS*110* family transposase |
| IS3 | 2615786~2617440 | 1655 | BTB_RS13165 | IS*4* like element IS*231*A family transposase |

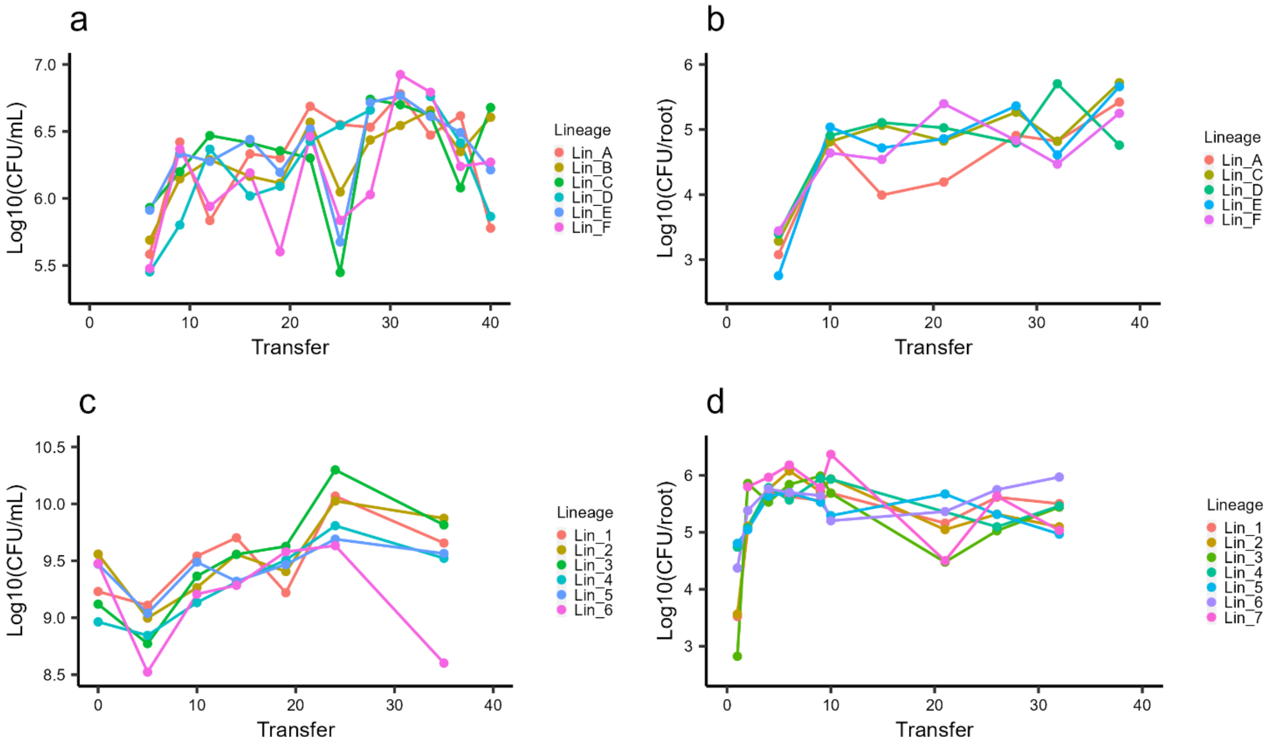


**Fig S1 Biofilm productivity of four experiments.** Data are from previous studies and replotted for better comparison

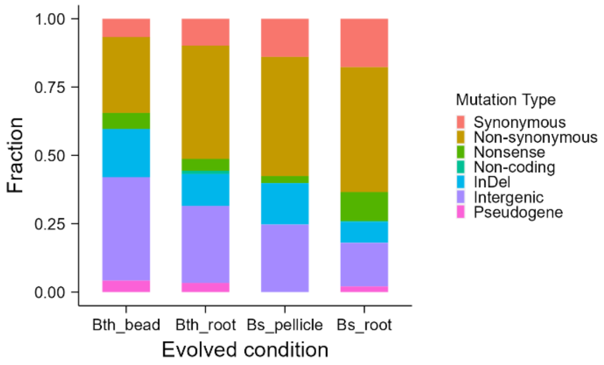


**Fig S2 Mutation spectrum.** Summary of mutational spectrum of the four adaptation models from all time points.

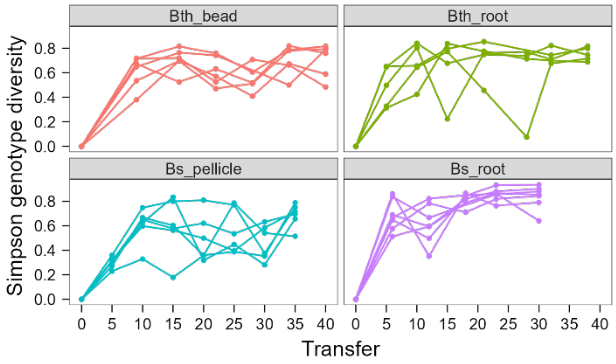


**Fig S3 Genotype diversity.** Dynamic distribution of genotype alpha diversity in each population of four adaptation models over time calculated using Simpson method.

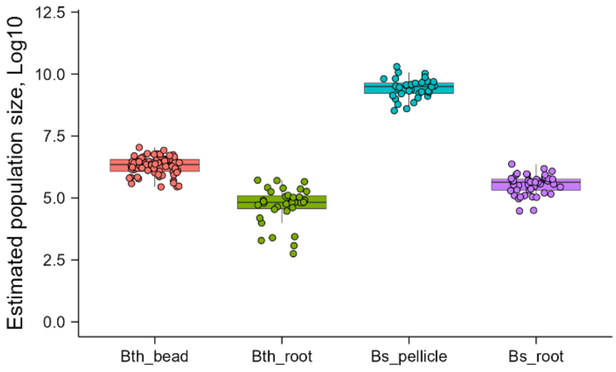


**Fig S5 Estimated population size of each population and timepoints calculated from biofilm productivity data.** Boxes indicate Q1–Q3, lines indicate the median, black circles filled with different color indicate the population size of each population.
