## Supplementary material for "Species and condition shape the mutational spectrum in experimentally evolved biofilms": Figure S4

### Supplementary Fig. 4A, Bth_bead lineage diagram

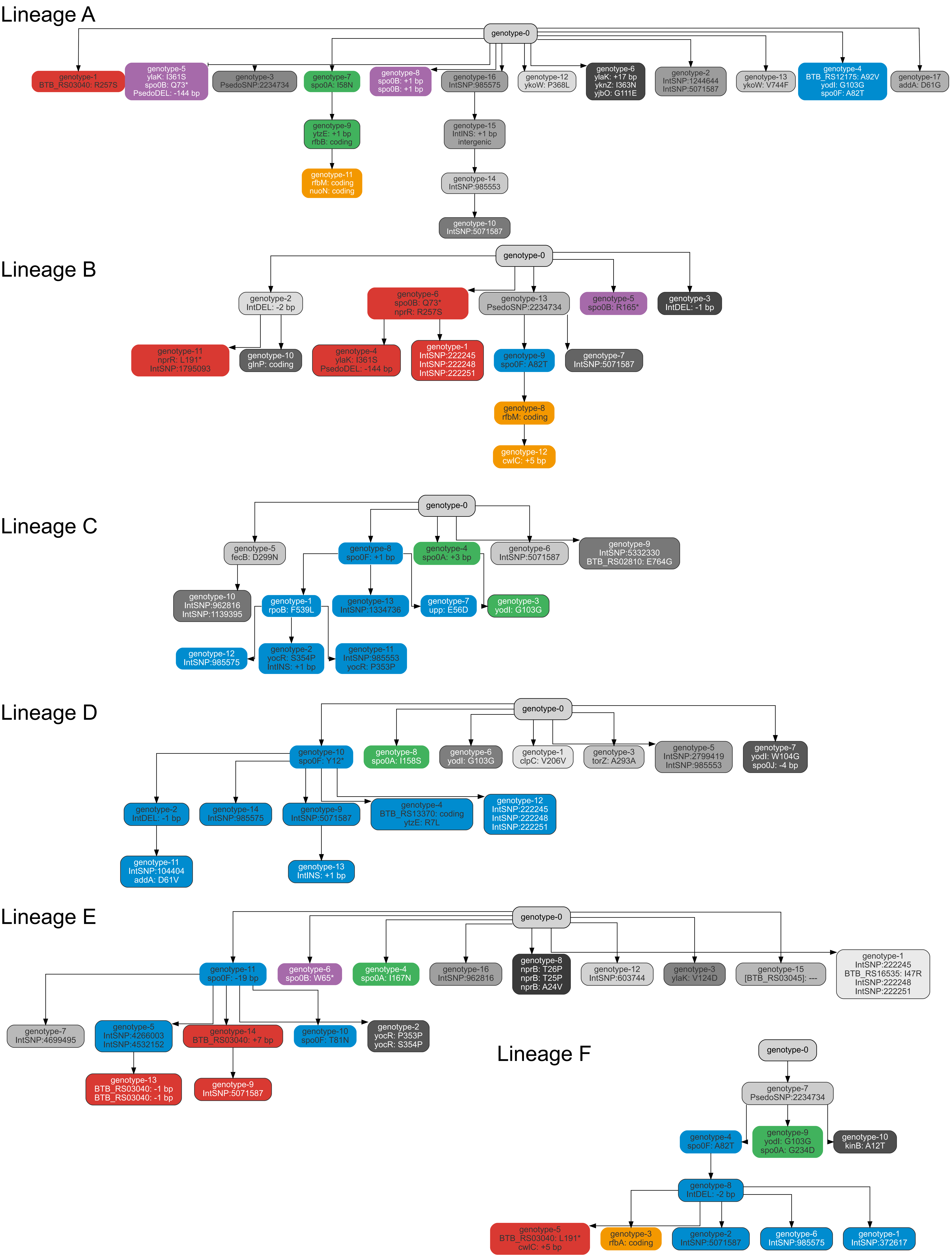

### Supplementary Fig. 4B, Bth_root lineage diagram

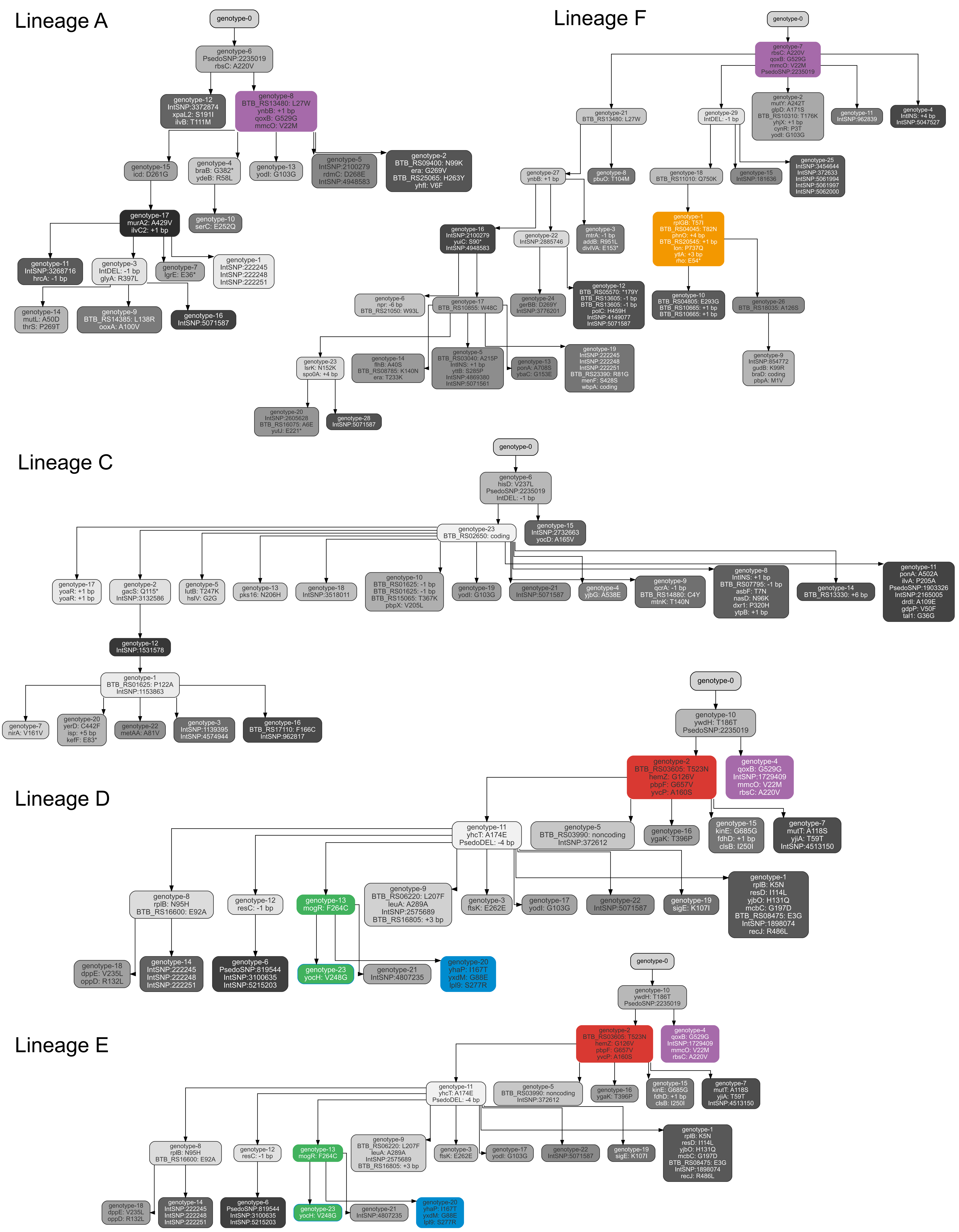

### Supplementary Fig. 4C, Bs_pellicle lineage diagram

#### Lineage 1

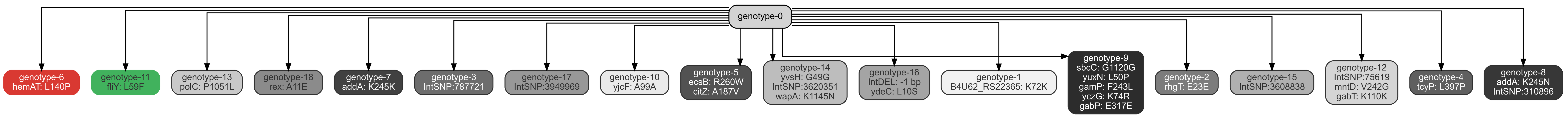


#### Lineage 2

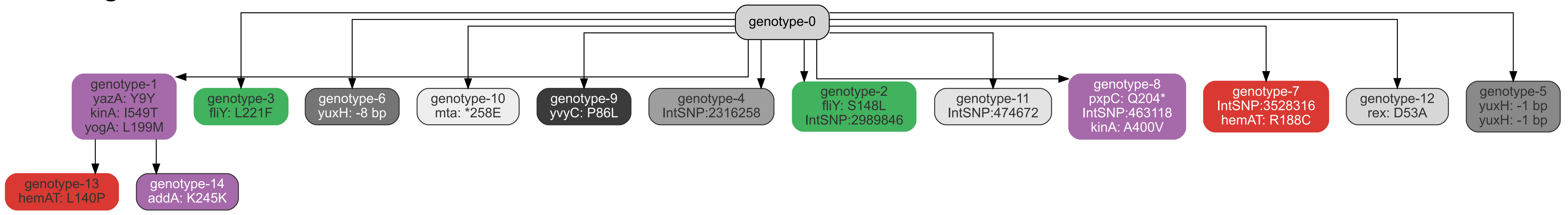


#### Lineage 3

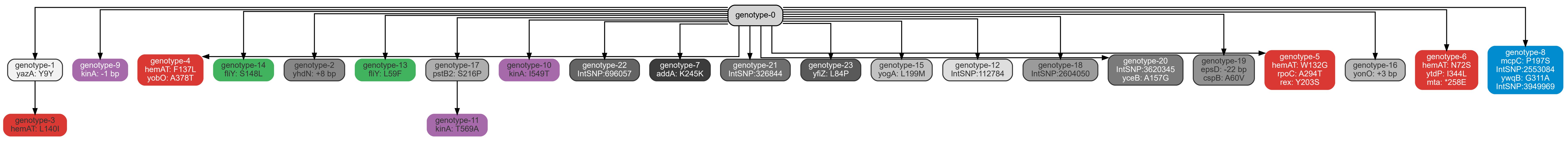


#### Lineage 4

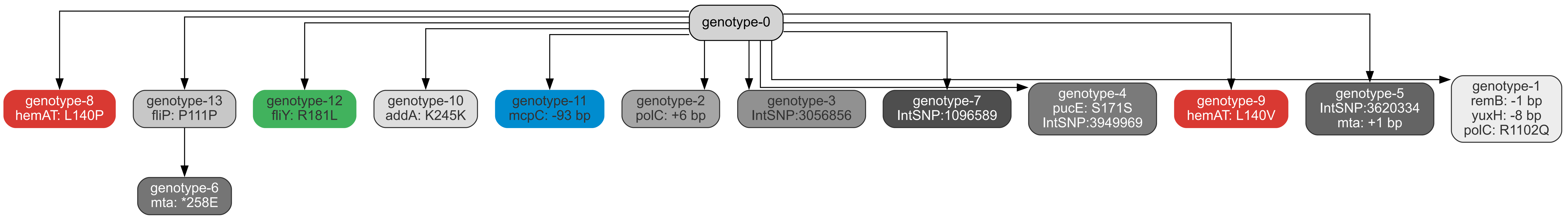


#### Lineage 5

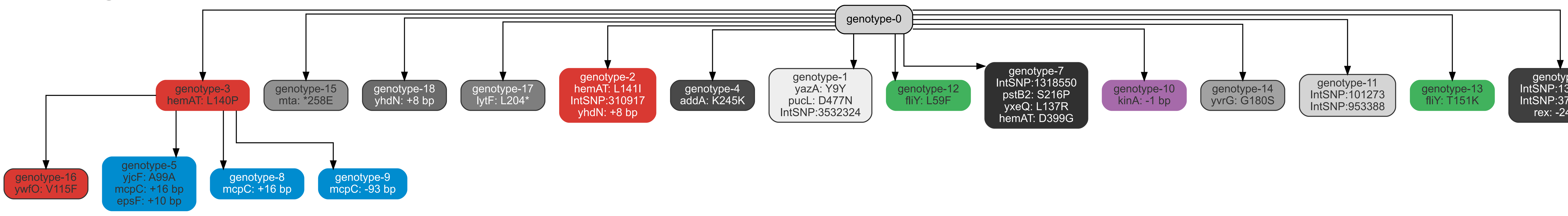


#### Lineage 6

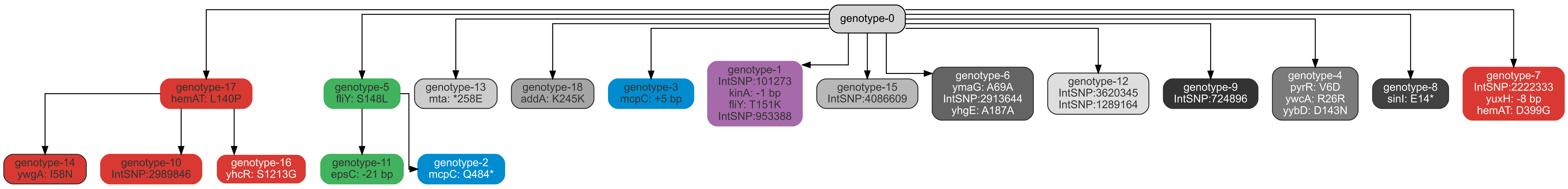

### Supplementary Fig. 4D, Bs_root lineage diagram

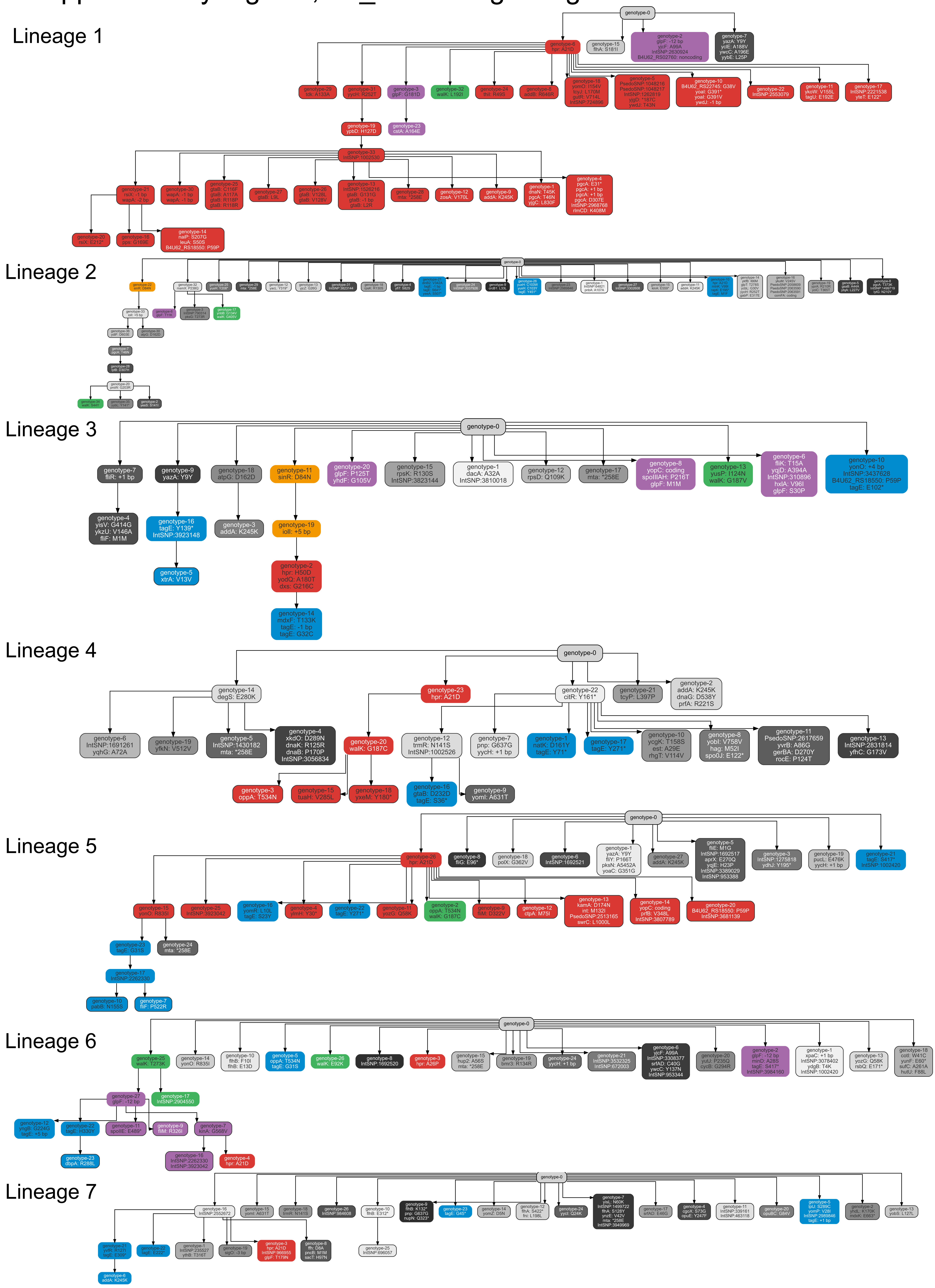
